# Binding Kinetics, Bias, Receptor Internalization and Effects on Insulin Secretion *in vitro* and *in vivo* of a Novel GLP-1R/GIPR Dual Agonist, HISHS-2001

**DOI:** 10.1101/2025.01.13.632834

**Authors:** Yusman Manchanda, Ben Jones, Gaelle Carrat, Zenouska Ramchunder, Piero Marchetti, Isabelle Leclerc, Rajamannar Thennati, Vinod Burade, Muthukumaran Natarajan, Pradeep Shahi, Alejandra Tomas, Guy A. Rutter

## Abstract

The use of incretin analogues has emerged in recent years as an effective approach to achieve both enhanced insulin secretion and weight loss in type 2 diabetes (T2D) patients. Agonists which bind and stimulate multiple receptors have shown particular promise. However, off target effects, including nausea and diarrhoea, remain a complication of using these agents, and modified versions with optimized pharmacological profiles and/or biased signaling at the cognate receptors are increasingly sought. Here, we describe the synthesis and properties of a molecule which binds to both glucagon-like peptide-1 (GLP-1) and glucose-dependent insulinotropic polypeptide (GIP) receptors (GLP-1R and GIPR) to enhance insulin secretion. HISHS-2001 shows increased affinity at the GLP-1R, as well as a tendency towards reduced internalization and recycling at this receptor *versus* FDA-approved dual GLP-1R/GIPR agonist tirzepatide. HISHS-2001 also displayed significantly greater bias towards cAMP generation *versus* β-arrestin 2 recruitment compared to tirzepatide. In contrast, G_αs_ recruitment was lower *versus* tirzepatide at the GLP-1R, but higher at the GIPR. Administered to obese hyperglycaemic *db/db* mice, HISHS-2001 increased circulating insulin whilst lowering body weight and HbA1c with similar efficacy to tirzepatide at substantially lower doses. Thus, HISHS-2001 represents a novel dual receptor agonist with an improved pharmacological profile.

## Introduction

Almost one third of North Americans are obese and two thirds are overweight^1^. Type 2 diabetes (T2D), driven in part by these increases in obesity in recent years, now affects almost one in ten of the population of westernized societies^2^, while the total number of cases worldwide is expected to rise to >600 million by 2045.

Originally identified as effective treatments for T2D^3,4^, GLP-1R agonists have proven effective in recent years in reducing both body weight and hyperglycaemia^5^. Dual and triple receptor agonists are proving even more effective^6,7^, with weight loss of 20% or more now achievable with the FDA-approved GLP1R-GIPR co-agonist, tirzepatide^8^. These agents bind to receptors in the endocrine pancreas, notably the pancreatic beta cell, potentiating the effects of glucose to stimulate insulin secretion, as well as to receptors in the brain, heart, kidney, adipose tissue and immune cells^9–11^ to promote energy expenditure and reduce appetite, amongst other beneficial effects. Dual incretin receptor agonists appear to deliver superior effects both through enhanced activation of insulin secretion^12^ but also centrally as a result of improved anti-emetic effects *versus* agonists which act through the GLP-1R alone^13^.

We have recently described a novel GLP-1R agonist, utreglutide (GL0034), with improved metabolic effects *versus* semaglutide^14^. Here, we sought to develop a similar, long-acting dual GLP-1R/GIPR receptor agonist and compare it to tirzepatide. We show that HISHS-2001 (GL0059) displays higher GLP-1R affinity and bias towards cAMP production over β-arrestin 2 recruitment, as well as comparable or improved metabolic effects in the *db/db* mouse model of diabetes *versus* tirzepatide. HISHS-2001 is thus a promising novel dual GLP-1R/GIPR agonist for trials in human obesity and T2D.

## Materials and Methods

### Animal maintenance and ethical approvals

Studies at Imperial College London were approved by the Animal Welfare and Ethical Review Body according to the UK Home Office Animals Scientific Procedures Act, 1986 (Project License PA03F7F0F to Isabelle Leclerc). All *in vivo* procedures at Sun Pharmaceuticals Inc. were approved by the Institutional Animal Ethics Committee (IAEC #608 and 650). Use of human islets was approved by the National Research Ethics Committee (NRES) London (Fulham), Research Ethics Committee no. 07/H0711/114, and by relevant national and local ethics committees including, where required, consent from next of kin. Human islet donor details are provided in Supplementary Table 1.

### Synthesis of HISHS-2001

Briefly, the parent peptide was synthesized by conventional solid-phase methods with Rink Amide as the starting resin used for synthesis. Following Fmoc de-protection of Rink Amide resin using Piperidine in DMF, Fmoc-Ser(tBu)OH coupling with Rink Amide was performed using DIPC and HOBt as coupling agent to yield Fmoc-Ser(tBu)-Resin. Uncoupled amino groups were capped after every amino acid coupling by acetylation. Next, selective de-blocking of the amino group of Fmoc-Ser(tBu)-Rink Amide Resin was performed using piperidine, followed by coupling with Fmoc-Pro-OH using HOBt and DIPC to yield Fmoc-Pro-Ser(tBu)-Rink Amide Resin. The above steps, i.e., selective capping, deblocking of Fmoc-protection of amino acid attached to the resin and coupling of next amino acid residue in sequence with Fmoc/Boc-protected amino group, were repeated for the remaining 37 amino acid residues. Once all amino acids are substituted on Rink Amide Resin, the 20th amino acid i.e., Lys was selectively deprotected and attached to the side chain. Finally, the crude compound was deprotected and cleaved from the resin using TFA/Ethanedithiol/TIPS mixture and purified through Preparative HPLC using Phenyl Silica column and Mobile phase (ammonium format buffer, aq. acetic acid, phosphate buffer). Fractions with purity above 98% (single major impurity limit NMT 0.5%) were pooled and desalted, and the desalted solution filtered (0.2 µm filter), and solvent distilled on rotavapor at 35-40°C under vacuum and lyophilised for ∼120 hours.

### Cell culture

HEK293 cells stably expressing human GLP-1R or human GIPR, each SNAP-tagged at their N-termini (Cisbio), were cultured in DMEM medium (Gibco), 10% FBS and 1% penicillin/streptomycin. DiscoverX GLP-1R-β-arrestin 2 cells were cultured in the manufacturer’s recommended medium.

Rat insulinoma INS-1 832/3 cell lines were cultured in RPMI 1640 medium (Gibco) supplemented with 10% FBS, 1% penicillin/streptomycin, 1 M HEPES buffer, 100 mM sodium pyruvate, and 0.05 mM β-mercaptoethanol. The cell lines used were INS-1 832/3 GLP-1R^-/-^ and INS-1 832/3 GIPR^-/-^ (kind gifts from Dr Jackeline Naylor, MedImmune/AstraZeneca^15^), and INS-1 832/3 stably expressing SNAP-tagged human GLP-1R (generated *in house* and previously described^16^).

Cell lines were kept in an incubator at a temperature of 37°C at a CO_2_ concentration of 5%.

### cAMP assays

HEK293-SNAP-GLP-1R, HEK293-SNAP-GIPR or GLP-1R-β-arrestin 2 cells were seeded overnight in 12-well plates. On the day of the assay, cells were resuspended and dispensed into white, 96-well plates containing the indicated concentration of agonist prepared in serum-free medium containing 0.1% BSA. After a 30-minute stimulation, cAMP detection reagents (cAMP dynamic kit, Cisbio) were added and the plate read 60 minutes later using an HTRF-compatible plate reader. Data were analysed using 3-parameter logistic fitting (GraphPad Prism 9.0) and normalized to the maximal fitted response of the reference ligand (GLP-1 or semaglutide) for presentation purposes.

### β-arrestin 2 recruitment assays

GLP-1R-β-arrestin 2 cells were seeded overnight in a 12-well plate. On the day of the assay, cells were resuspended and dispensed into white, 96-well plates containing the indicated concentration of agonist prepared in serum-free medium containing 0.1% BSA. After a 30-minute stimulation, β-arrestin detection reagents (DiscoverX) were added and the plate read 60 minutes later by luminescence. Data were analyzed using 3-parameter logistic fitting (GraphPad Prism 9.0) and normalised to the maximal fitted semaglutide for presentation purposes.

### Calculation of bias

Bias between cAMP and β-arrestin 2 was determined using a modified operational model of agonism. Concentration response data was fitted with previously described equations^17^ to derive transduction ratios (τ/KA) for each agonist and pathway. Log (τ/KA) values were normalized by subtracting log (τ/KA) for semaglutide in each pathway, giving Δlog (τ/KA), and to determine the bias between the two pathways, Δlog (τ/KA) values were subtracted, yielding ΔΔlog (τ/KA). This value is referred to as “bias factor” in the text.

### GLP-1R binding kinetics

HEK293-SNAP-GLP-1R cells were seeded overnight in 12-well plates. On the day of the assay, cells were labelled with 40 nM Lumi4-Tb (Cisbio) in complete medium for 30 minutes. After washing, labelled cells were preincubated for 20 minutes in HBSS supplemented with 0.1% BSA and a metabolic inhibitor cocktail (10 mM NaN_3_, 20 mM 2-deoxyglucose) to inhibit GLP-1R endocytosis that would otherwise influence binding assays. Ligand binding was then detected in real time using TR-FRET using a Flexstation 3 plate reader (Molecular Devices). Ligand conditions included four or more concentrations of unlabelled HISHS-2021 or tirzepatide in combination with 10 nM exendin(9-39)-FITC, and exendin(9-39)-FITC alone at four concentrations to determine its own kinetic parameters. Plate reader settings included 340 nm λexcitation, 520 nm (cut-off 495 nm) and 620 nm (cut-off 570 nm) λemission, 400 μsec delay and 1500 μsec integration time. Association and dissociation rate constants were calculated using the kinetics of competitive binding algorithm in GraphPad Prism 9.0.

### G_αs_ recruitment by NanoBiT complementation assay

INS-1 832/3 GLP-1R^-/-^ and GIPR^-/-^ cells were seeded onto 6-cm dishes the day before transfection and transfected the following day using Lipofectamine 2000 (Thermo Fisher Scientific) with 1 µg plasmid DNA to 2 µL Lipofectamine 2000 ratios. Cells were transfected with 1.7 µg GLP-1R-SmBiT or 1.7 µg GIPR-SmBiT (cloned *in house*) and 1.7 µg G_αs_-LgBiT (gift from Prof Nevin Lambert, Medical College of Georgia, USA). 4 hours prior to the assay, the cell media was replaced with RPMI supplemented with 3 mM glucose. Cells were then washed with PBS, detached with ethylenediaminetetraacetic acid (EDTA), and resuspended in HBSS buffer, after which Nano-Glo® LCS Dilution Buffer and Nano-Glo® LCS Live Cell Substrate were added, according to the manufacturer’s protocol (Promega), and a baseline reading was taken for 8 minutes. Cells were subsequently stimulated with the indicated concentration of agonist for 30 minutes with a reading taken every 30 sec. The luminescence (RLU) measured for each agonist was normalised to the baseline reading and to the vehicle luminescence reading. The AUCs were calculated and a dose-response curve generated in GraphPad Prism 9.0.

### Lipid Nanodomain isolation by membrane fractionation

GLP-1R recruitment into lipid nanodomains was measured using INS-1 832/3 SNAP-GLP-1R cells as follows: 2×10^6^ cells were seeded on 6-cm dishes the day before fractionation. On the day of fractionation, cells were stimulated with 100 nM agonist for 2 minutes, washed with ice-cold PBS and osmotically lysed on ice in lysis buffer [20 mM Tris-HCl, pH 7, 1% protease inhibitor (Roche) and 1% phosphatase inhibitor (Sigma-Aldrich)]. Lysates were passed through 21 gauge needles and ultracentrifuged at 41,000 rpm at 4°C for 1 hour. Pellets were resuspended in PBS supplemented with 1% Triton-X100, 1% protease inhibitor and 1% phosphatase inhibitor, incubated under rotation at 4°C for 30 minutes and ultracentrifuged again at 41,000 rpm at 4°C for 1 hour. The detergent-soluble membrane fractions (DSM, supernatants) were removed, and the detergent-resistant membrane fractions (DRM, pellets) resuspended in 1% SDS with 1% protease inhibitor and 1% phosphatase inhibitors, sonicated, centrifuged at 1,300 rpm at 4°C for 5 minutes and stored at -20°C for Western blot analysis. The DRMs and DSMs were diluted 1:1 with 2x TBE urea buffer (200 mM Tris-HCl, pH 6.8, 5% w/v SDS, 8 M urea, 100 mM DTT and 0.02% w/v bromophenol blue), incubated at 37°C for 10 minutes and resolved by SDS–polyacrylamide gel electrophoresis (10% acrylamide gels). Protein transfer to PVDF membranes (Immobilon-P, 0.45-μm pore size, IPVH00010, Merck) was achieved using a wet transfer system (Bio-Rad). Blotted membranes were blocked in 5% skimmed milk in TBS-Tween buffer, incubated with primary rabbit anti-SNAP monoclonal antibody (1:1,000, New England Biolabs), followed by secondary HRP-conjugated anti-rabbit antibody (1:2,000, Invitrogen) in the same buffer and developed using the Clarity Western enhanced chemiluminescence substrate system (1705060, Bio-Rad) in a Xograph Compact X5 processor. Membranes were then stripped in stripping buffer (20 g SDS, 7.6 g Trizma-Base, pH 6, supplemented with 1% β-mercaptoethanol) at 50°C for 15 minutes, blocked as above, and incubated with rat primary anti-flotillin antibody (2.5 μg/μL, BioLegend), followed by secondary HRP-conjugated anti-rat antibody (1:2,500, Sigma-Aldrich). Specific band densities were quantified using Fiji ImageJ v1.53c. SNAP bands were normalised to flotillin and expressed as fold-change over vehicle.

### GLP-1R internalization and recycling assays

#### GLP-1R internalization

INS-1 832/3 SNAP-GLP-1R cells were seeded in 14-mm microwell glass bottom MatTek dishes (MatTek Life Sciences). Prior to imaging, the cells were labelled with SNAP-Surface® 549 (New England Biolabs) for 20 minutes in full media. Cells were then washed and imaged in Live Cell Imaging Solution (Thermo Fisher Scientific) by time-lapse spinning disk microscopy using a Nikon Eclipse Ti spinning disk microscope with an ORCA-Flash 4.0 camera (Hamamatsu) and Metamorph software (Molecular Devices) with a 60x/1.4 NA oil objective 1 minute before and 9 minutes post-stimulation with 100 nM agonist, with images acquired at 6 sec intervals. Time-lapse images were analysed in Fiji ImageJ v1.53c using a macro designed by Imperial College Facility for Light Microscopy (FILM). Loss of receptor from the plasma membrane was quantified per frame by calculating full width at half maximum (FWHM) values across the cell membrane and normalized to the average membrane receptor at baseline, and results were expressed as a percentage of internalized receptor over time.

#### GLP-1R recycling

INS-1 832/3 SNAP-GLP-1R cells were seeded in 96-well imaging plates. Cells were washed twice with PBS and stimulated with 100 nM of agonist in complete medium for 1 hour to induce receptor internalization, washed twice with PBS and labelled with 100 nM exendin-4-TMR for 1 hour or 3 hours to detect receptor reappearance at the cell surface, followed by a wash in PBS and imaging on a high content Nikon Eclipse Ti2 microscope with a 20x/0.5 NA objective lens, 9 fields of view were acquired per well and analysed in Fiji ImageJ v1.53c. Images were phase segmented, and fluorescence images were background corrected, and fluorescent intensities measured from segmented cell-containing regions, with measurements normalised to signal from exendin-4-TMR-negative wells.

### *Ex vivo* Ca^2+^ mobilisation in mouse islets

Imaging of intact islets 24 hours post-isolation was performed essentially as described^18,19^. In brief, islets from individual wildtype mice were preincubated for 1 hour in Krebs-Ringer bicarbonate–Hepes (KRBH) buffer (140 mM NaCl, 3.6 mM KCl, 1.5 mM CaCl_2_, 0.5 mM MgSO_4_, 0.5 mM NaH_2_PO_4_, 2 mM NaHCO_3_, 10 mM Hepes, saturated with 95% O_2_/5% CO_2_; pH 7.4) containing 0.1% (w/v) bovine serum albumin (BSA), 6 mM glucose (KRBH G6), and the Ca^2+^-responsive dye Cal-520 AM (2 µM, AAT Bioquest). Treatments with KRBH G6 ± 100 nM agonist, 11 mM glucose, or 20 mM KCl were manually added to the islet dishes by pipetting at the indicated time points. To ensure that the islets remained immobile, these were pre-encased into Matrigel (356231, Corning) and imaged at 37°C on glass-bottom dishes (P35G-1.5-10-C, MatTek Life Sciences). Imaging was performed using a using a Nikon Eclipse Ti spinning disk microscope with an ORCA-Flash 4.0 camera (Hamamatsu) and Metamorph software (Molecular Devices) with the following settings: lexcitation 488, lemision 510 nm), 20x/0.5 NA air objective. Raw fluorescence intensity traces from whole-islet ROIs were extracted using Fiji ImageJ v1.53c. Responses were plotted relative to the average fluorescence intensity per islet during the KRBH G6 baseline period, before agonist addition.

### Islet insulin secretion

Secretion assays were performed in purified mouse and human islets. Human islets from the European Consortium for Islet Transplantation were used for these experiments. Secreted and total insulin were measured essentially as per ^14^. In brief, islets (10/well) were incubated in triplicate for each condition and treatment. Islets (human or mouse) were pre-incubated for 1 hour in KRBH buffer containing 1% (w/v) BSA and 3 mM glucose before incubation with 11 mM glucose ± 10 nM agonist in KRBH in a shaking 37°C water bath (80 rpm) for 1 hour. Following stimulation, supernatants containing the secreted insulin were collected, centrifuged at 1000 g for 5 minutes, and transferred to fresh tubes. To determine total insulin contents, islets were lysed using acidic ethanol [75% (v/v) ethanol and 1.5 mM HCl]. The lysates were sonicated 3 × 10 sec in a water bath and centrifuged at 10,000 g for 10 minutes, and the supernatants collected. The samples were stored at −20°C until the insulin concentration was determined using an Insulin Ultra-Sensitive HTRF Assay kit (62IN2PEG, Cisbio) according to the manufacturer’s instructions. GraphPad Prism 9.0 was used for the generation of the standard curve and sample concentration extrapolation. The total insulin content was calculated by adding the secreted insulin to the insulin content of the lysates.

### Treatment of *db/db* mice

*Db/db* mice [C57BL/KsJ-db/db male/female mice (age 8-11 weeks; body weight 40-60 g; n=8) were randomised based on HbA1c levels. The mice, which were procured from Laboratory Animal Resources (LAR; Sun Pharma Advanced Research Company Ltd.), were housed in individual ventilated cages with free access to food and water and maintained on a 12-hour light/dark cycle. Animals were acclimatized for 3 days. On day 0, each animal was weighed using a digital weighing balance. Blood was collected by retro-orbital plexus puncture. Blood glucose level was measured with glucose strips using Blood Glucose Meter (One TouchTM UltraTM; LIFESCAN, Johnson & Johnson) and % HbA1c was measured using A1C Now+® (PTS Diagnostics). Vehicle (placebo), HISHS-2001 (4.5, 9 and 18 nmol/kg), or tirzepatide (180 nmol/kg) were injected subcutaneously in the neck region of the animals on every third day for 4 weeks (q3d*10). Body weight was monitored. On day 28 of the study, 24 hours following the last dose, blood was collected for measurement of HbA1c, triglycerides (Colorimetric Assay), and insulin (ELISA).

### Food consumption

Animals were house at 4 mice per cage with pre-weighed quantities of rodent diet. The amount of food left was measured serially. At the end of the 28 days treatment period, the amount of diet consumed per cage was determined and represented as food consumption per group.

### Pharmacokinetic analysis

Plasma pharmacokinetic study of HISHS-2001 and tirzepatide was performed in male CD-1 mice. 90 animals were procured from Laboratory Animal Resources (LAR). Animals were acclimatized for one day. On day 0, each animal was weighed. Animals were divided into 18 groups (9 groups for HISHS-2001 and 9 groups for tirzepatide), each containing 5 male animals and representing one time point. Animals were injected subcutaneously at 1 mg/kg for the corresponding agonist (dose volume: 10 mL/kg) using 26½ gauge needles. At 1, 2, 4, 8, 12, 24, 48, 72 and 96 hours, ∼500 µL of blood was collected by retro-orbital plexus puncture using capillary micro-centrifuge tubes containing anticoagulant (15 µL 10% K_2_EDTA). Plasma was separated from blood by centrifugation at ∼3,300 rpm for 10 minutes at 4°C. HISHS-2001 concentrations were estimated using liquid chromatography with tandem mass spectrometry (LC-MS/MS). Lower Limit of Quantification (LLOQ) was 2.14 ng/mL for HISHS-2001 and 2.04 ng/mL for tirzepatide.

### Statistics

Data analysis was performed using GraphPad Prism 9.0, involving ANOVA with appropriate post-hoc correction for multiple testing, and p<0.05 was considered significant.

## Results

### Receptor pharmacology of HISHS-2001 resembles that of tirzepatide

The structure of HISHS-2001 is shown in Figure 1A. Similarly to tirzepatide, it consists of a 39 amino acid peptide backbone modified with a fatty acid side chain in Lys 20 and features two α-amino isobutyric acid (Aib) residues in positions 2 and 13. The main difference with tirzepatide lies on the composition of its side chain with a distinct linker sequence that couples the peptide to an albumin-binding fatty acid moiety.

**Figure 1.**
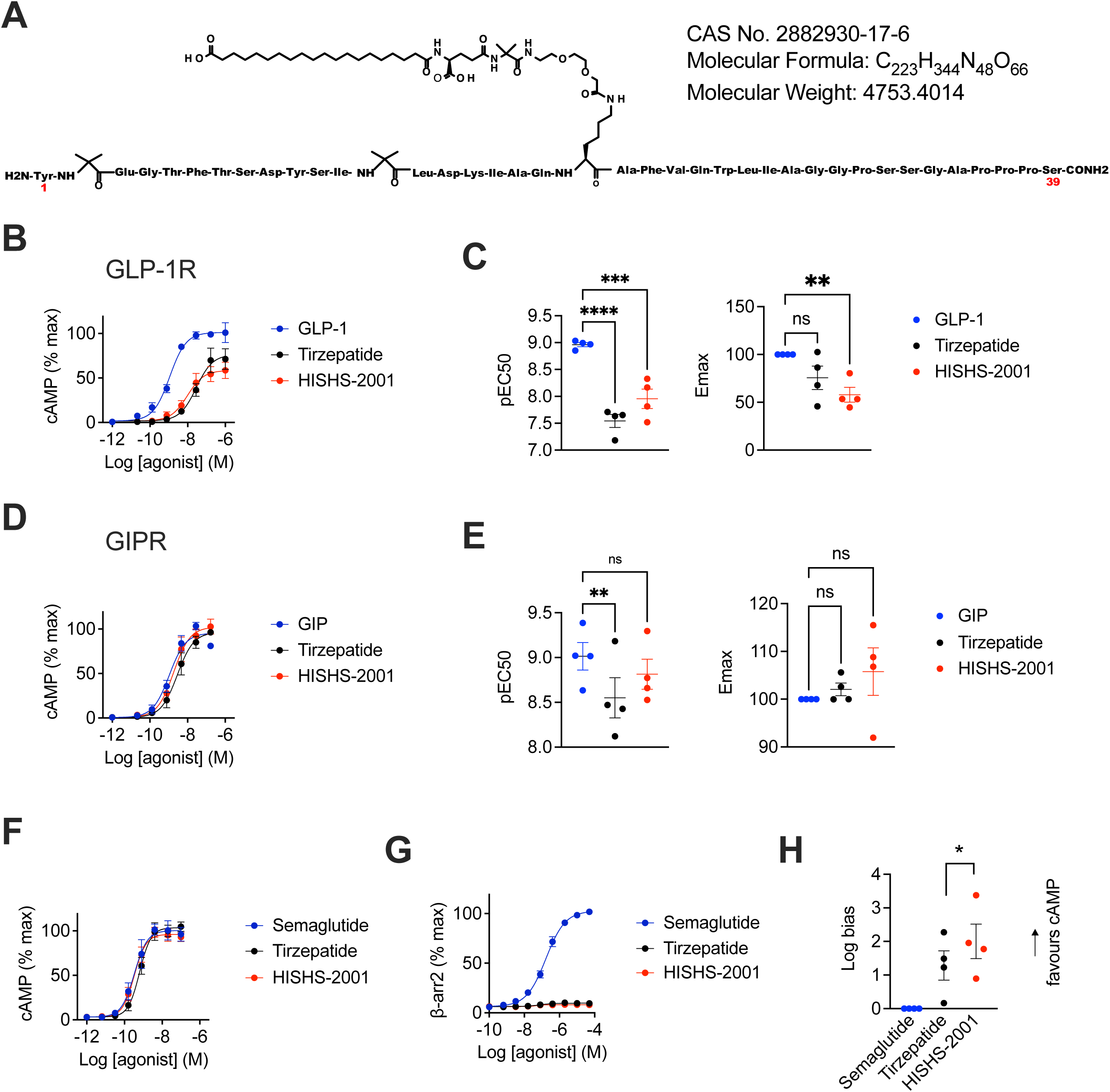
Pharmacological characterisation of HISHS-2001 at GLP-1R and GIPR. (**A**) Schematic showing HISHS-2001 sequence and other formula details. (**B**) cAMP dose responses for the indicated agonists in HEK293-SNAP-GLP-1R cells, n=5. (**C**) Potency (pEC50) and maximal response estimates from (B). (**D**) cAMP dose responses for the indicated agonists in HEK293-SNAP-GIPR cells, n=4. (**E**) Potency (pEC50) and maximal response estimates from (D). (**F**) cAMP responses in DiscoverX GLP-1R-β-arrestin 2 cells, n=4. (**G**) β-arrestin 2 recruitment responses in DiscoverX GLP-1R-β-arrestin 2 cells, n=4. (**H**) Bias factor calculated using from transduction ratios (*τ*/*K*_A_) method for HISHS-2021 and tirzepatide compared to semaglutide using data from (F) and (G). Data is shown as mean +/- SEM; ns, non-significant; *p<0.05; **p<0.01; ***p<0.001; ****p<0.0001 by one-way ANOVA with Dunnett’s post-hoc test.

To explore the biological actions of this molecule, we first examined agonist-induced cAMP changes in HEK293 cells expressing either SNAP-tagged human GLP-1R or GIPR. Both HISHS-2001 and tirzepatide displayed significantly reduced potency compared to native GLP-1(7-36)NH_2_ at the GLP-1R, although HISHS-2001 appeared to be the more potent of the two synthetic ligands (Figure 1B, C). HISHS-2001 and tirzepatide both behaved as partial agonists in this system, in keeping with the previously described pharmacology of tirzepatide, which shows reduced maximal ability to activate G_αs_ signalling^20^. Both synthetic agonists showed similar potencies to native GIP(1-42) in GIPR-expressing HEK293 cells, although, again, tirzepatide was the least potent of all the agonists at the GIPR (Figure 1D, E).

As tirzepatide is reported to be a biased agonist with minimal β-arrestin recruitment at the GLP-1R, we next aimed to assess signal bias for both agonists using PathHunter® cells, which allow cAMP and β-arrestin 2 responses to be determined in parallel. In this cell model, both HISHS-2001 and tirzepatide showed similar potency to semaglutide for cAMP production at the GLP-1R, although HISHS-2001 was moderately more potent (Figure 1F); however, both HISHS-2001 and tirzepatide showed profoundly reduced ability to drive β-arrestin 2 recruitment (Figure 1G). Calculation of bias from these data revealed both dual agonists were strongly biased towards cAMP signalling, with HISHS-2001 appearing to be the most strongly biased of the two (Figure 1H).

Some reports indicate that G protein-biased GLP-1R agonists tend to show lower affinity as a result of faster receptor dissociation kinetics^14,21^. We therefore used a Förster resonance energy transfer (FRET)-based assay^14^ to determine the binding affinities of each ligand at the GLP-1R through competitive kinetics with the fluorescent GLP-1R ligand exendin(9-39)-FITC (Figure 2). K_on_ values for HISHS-2001 and tirzepatide were similar (Figure 2A), but K_off_ (Figure 2B) was significantly lower for HISHS-2001 than for tirzepatide, such that the calculated K_d_ value (Figure 2C) was also lower. Thus, HISHS-2001 displays tighter binding at the GLP-1R than tirzepatide, which could explain its modestly higher potency for cAMP production.

**Figure 2.**
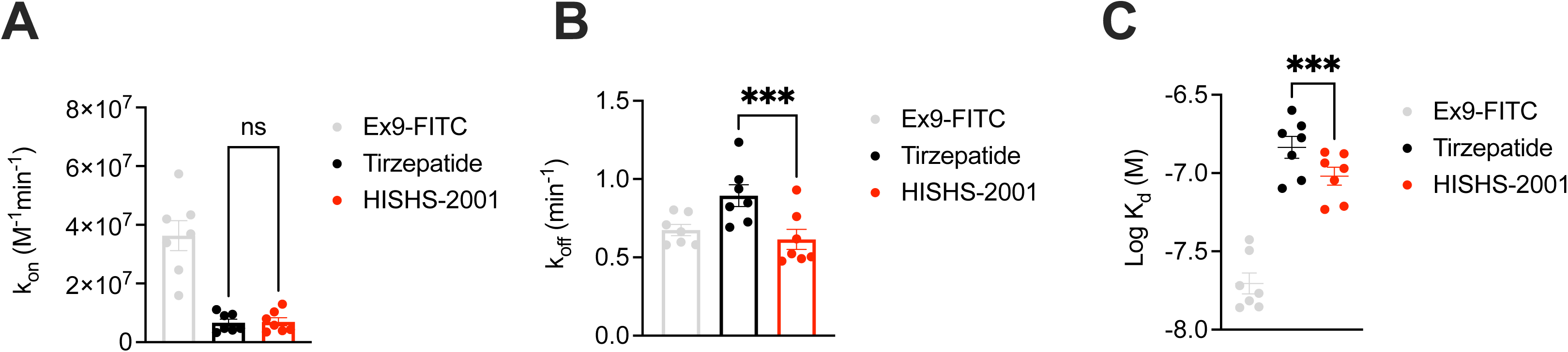
HISHS-2001 *versus* tirzepatide GLP-1R binding kinetics. (**A**) K_on_ calculated from competitive kinetics assay in HEK293-SNAP-GLP-1R cells, n=4. (**B**) K_off_ calculated from competitive kinetics assay in HEK293-SNAP-GLP-1R cells, n=4. (**C**) Calculated affinity from data shown in (A) and (B). Data is shown as mean +/- SEM; ns, non-significant; ***p<0.001 by paired t-test.

### Subtle changes in G_αs_ coupling triggered by HISHS-2001 compared to tirzepatide in pancreatic beta cells

A NanoBiT complementation assay was next used in INS-1 832/3 rat pancreatic beta cells to explore the coupling between each receptor and G_αs_ with both synthetic agonists. Whilst HISHS-2001 triggered a lower degree of G_αs_ recruitment *versus* tirzepatide at the GLP-1R (Figure 3A, B), a tendency was observed towards more efficient coupling at the GIPR with this agonist (Figure 3C, D).

**Figure 3.**
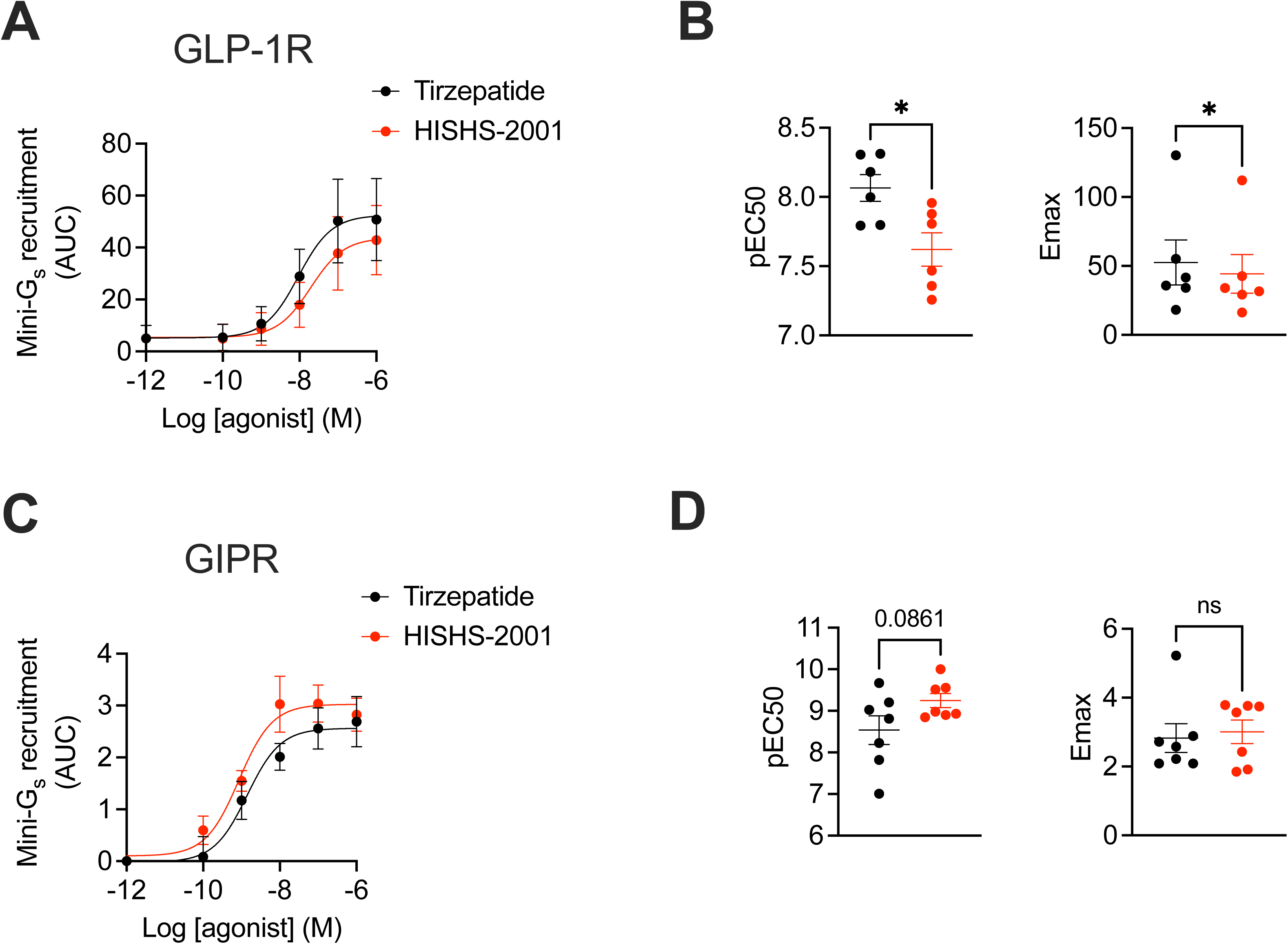
HISHS-2001 *versus* tirzepatide G_αs_ protein coupling at the GLP-1R and GIPR in beta cells. (**A**) GLP-1R G_αs_ recruitment dose response to HISHS-2001 and tirzepatide at indicated doses by NanoBiT complementation in INS-1 832-3 GLP-1R^-/-^ cells transfected with GLP-1R-SmBiT and G_αs_-LgBiT. (**B**) Potency (pEC50) and maximal response from (A); n=6. (**C**) GIPR G_αs_ recruitment dose response to HISHS-2001 and tirzepatide at indicated doses by NanoBiT complementation in INS-1 832-3 GIPR^-/-^ cells transfected with GIPR-SmBiT and G_αs_-LgBiT. (**D**) Potency (pEC50) and maximal response from (C); n=7. Data is shown as mean +/- SEM; ns, non-significant; *p<0.05 by paired t-test.

### HISHS-2001 and tirzepatide elicit a similar GLP-1R trafficking profile and cholesterol-rich lipid nanodomain partitioning of the receptor

GLP-1R internalization, assessed by confocal imaging of SNAP-tagged human GLP-1R stably expressed in INS-1 832/3 cells, revealed a slower endocytosis of the receptor following stimulation with both tirzepatide and HISHS-2001 when compared to the non-biased GLP-1R agonist semaglutide (Figure 4A, B), as previously observed for tirzepatide^22^. A faster GLP-1R recycling for both receptors *versus* semaglutide was also observed, although this only reached significance with tirzepatide (Figure 4C).

**Figure 4.**
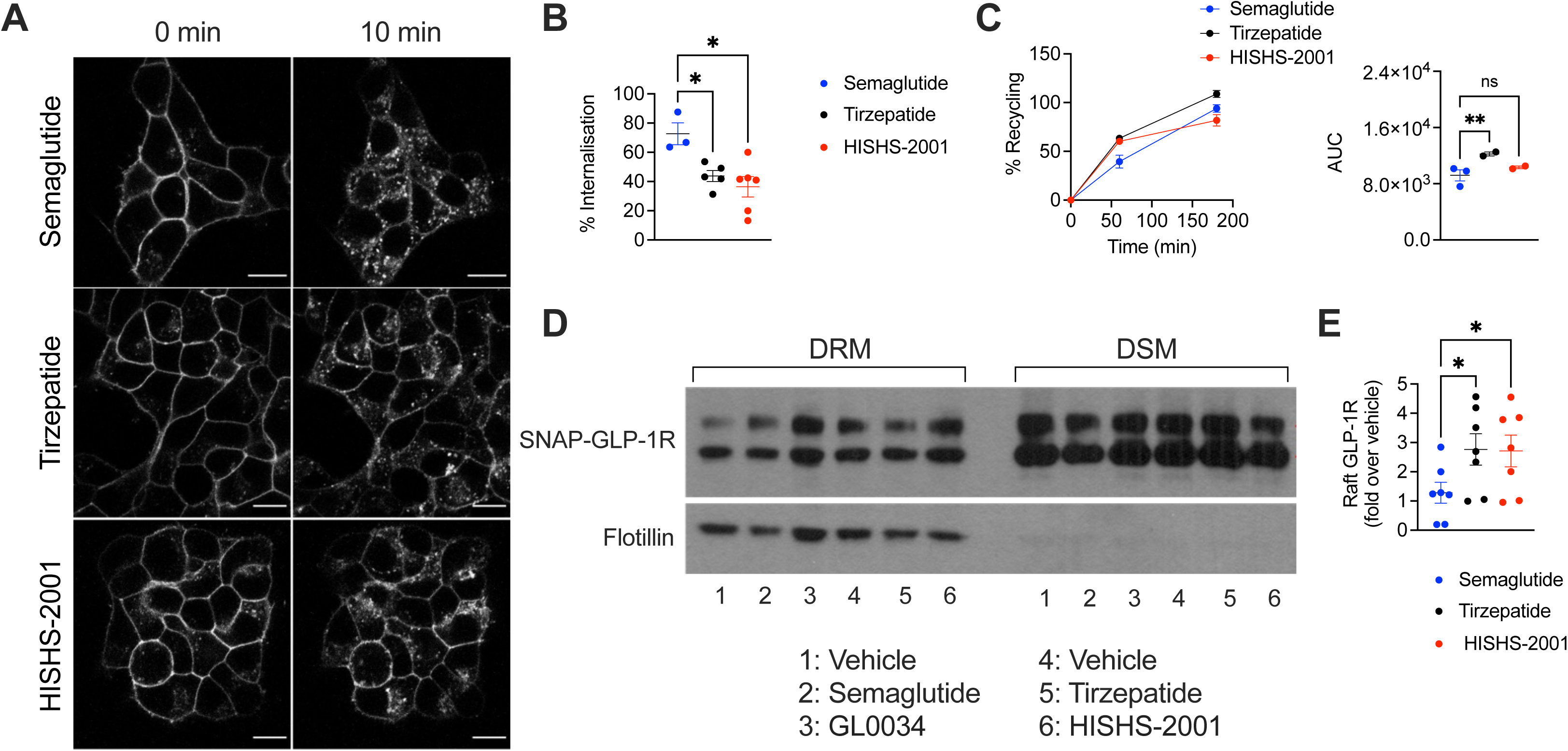
GLP-1R trafficking profiles of HISHS-2001 *versus* tirzepatide or semaglutide in beta cells. (**A**) Representative images of SNAP-GLP-1R subcellular localisation at 0- and 10-minutes post-stimulation with 100 nM HISHS-2001, tirzepatide, or semaglutide in INS-1 832/3 SNAP-GLP-1R cells. (**B**) Percentage of SNAP-GLP-1R internalization with the indicated agonist calculated from (A); n=3-7. (**C**) Percentage of SNAP-GLP-1R recycling to the plasma membrane in INS-1 832/3 SNAP-GLP-1R cells in response to 100 nM HISHS-2001, tirzepatide, or semaglutide, with corresponding AUCs shown; n=2-3. (**D**) Cholesterol-rich lipid nanodomain segregation of SNAP-GLP-1R in INS-1 832/3 SNAP-GLP-1R cells under vehicle conditions or in response to 100 nM HISHS-2001, tirzepatide, semaglutide, or semaglutide analogue GL0034^14^; DRM, detergent-resistant membrane fractions; DMS, detergent-soluble membrane fractions; flotillin indicates cholesterol-rich lipid nanodomain enrichment. (**E**) Quantification of SNAP-GLP-1R/flotillin from (D); n=7. Data is shown as mean +/- SEM; ns, non-significant; *p<0.05; **p<0.01 by one-way ANOVA with Dunnett’s post-hoc test.

GLP-1R signalling efficacy is impacted by recruitment of the receptor to cholesterol-rich lipid nanodomains^23^. After exposure to agonists, we employed a biochemical fractionation approach to separate detergent-soluble and -resistant plasma membrane fractions in INS-1 832/3 beta cells stably expressing SNAP-tagged human GLP-1R. Western blot analyses of these fractions showed that both tirzepatide and HISHS-2001 drove similarly elevated incorporation of GLP-1R into lipid rafts *versus* semaglutide, with no differences between the two dual agonists (Figure 4D, E).

### HISHS-2001 and tirzepatide elicit similar levels of Ca^2+^ dynamics and potentiation of insulin secretion in primary mouse and human islets

We next explored the degree of signal transduction elicited by both synthetic dual agonists in primary islets. We first analyzed their capacity to elicit intracellular Ca^2+^ rises with the intracellular fluorescent dye Cal520 in islets purified from wildtype mice. Here, both tirzepatide and HISHS-2001 caused a similar potentiation of glucose-induced intracellular Ca^2+^ increases, with tirzepatide tending to be more effective at 6 mM glucose, and HISHS-2001 at 11 mM glucose (Figure 5A, B). We next examined the capacity of both agonists to potentiate glucose-stimulated insulin secretion in both mouse and human islets, observing a similar potentiating effect by the dual receptor agonists as that promoted by semaglutide at 11 mM glucose in islets from both species (Figure 5C, D).

**Figure 5.**
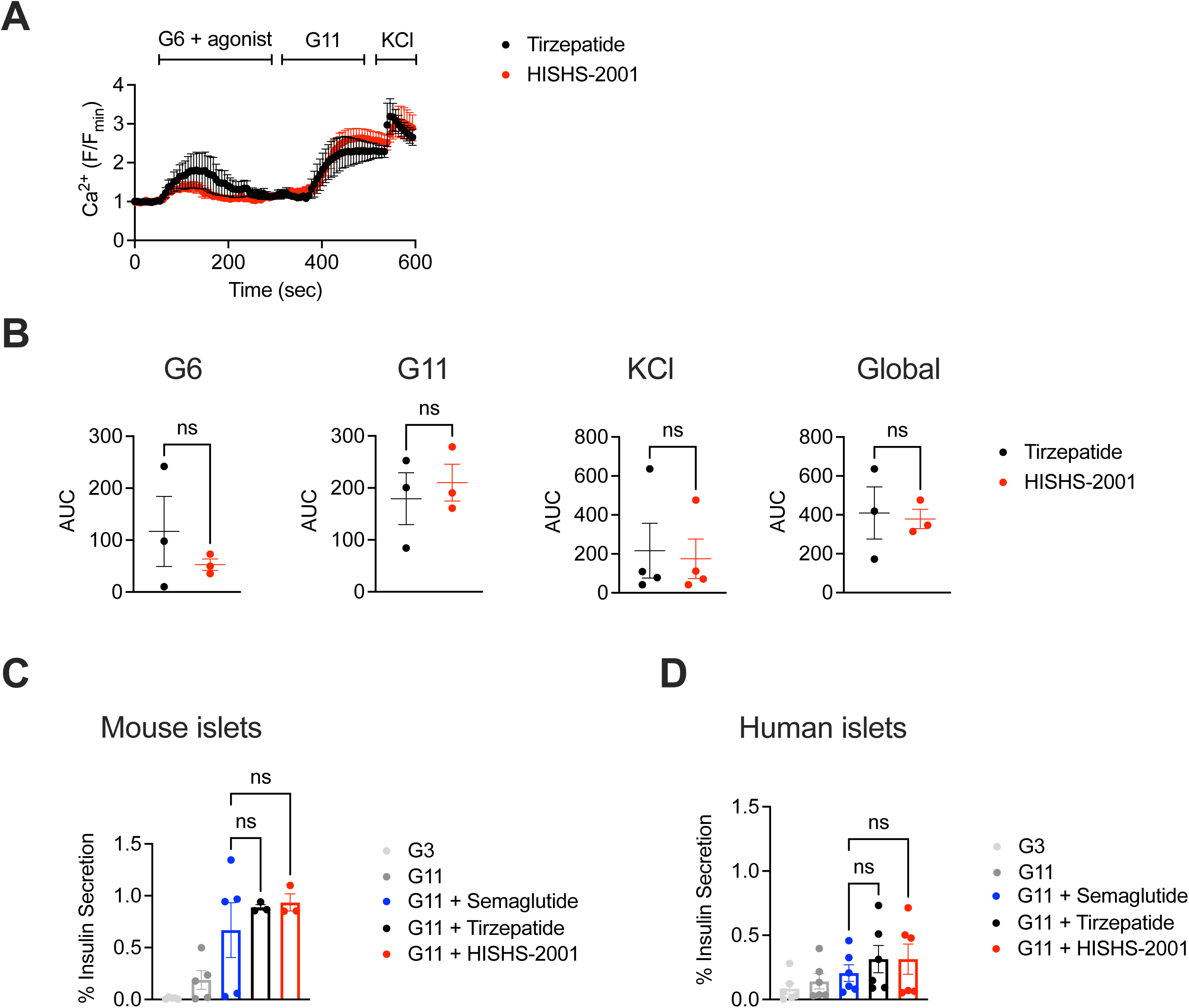
*Ex vivo* HISHS-2001 *versus* tirzepatide downstream functional responses in primary islets. (**A**) *Ex vivo* intracellular Ca^2+^ increases in wildtype mouse islets in response to 100 nM HISHS-2001 or tirzepatide, shown as F/Fo (Fo, average baseline signal); G6, 6 mM glucose; G11, 11 mM glucose; KCl, 20 mM. (**B**) AUCs calculated for the G6, G11, KCl and global responses; n=3. (**C**, **D**) *Ex vivo* insulin secretion from mouse (C) and human (D) islets at indicated glucose concentrations and in response to 100 nM HISHS-2001, tirzepatide, or semaglutide; G3, 3 mM glucose; G11, 11 mM glucose; n=3-6. Data is shown as mean +/- SEM; ns, non-significant by paired t-test or one-way ANOVA with Dunnett’s post-hoc test.

### Effects of HISHS-2001 on glucose homeostasis *in vivo*

Finally, a chronic *in vivo* study was performed where obese mixed sex *db/db* mice were exposed to HISHS-2001 or tirzepatide for 28 days. The stability of each drug in the circulation was similar (Supplementary Table 2). We also observed similar reductions in circulating triglycerides and body weight with both agonists (Figure 6A, B), but with HISHS-2001 used at 10 times lower dose compared to tirzepatide. The effects of each drug on food intake were similar (∼50 % lowering; Supplementary Table 3), and both agonists also caused similar reductions in HbA1c with HISHS-2001 used at 10-fold reduced dose. Nevertheless, increases in circulating insulin were significantly greater for HISHS-2001 despite the reduced dosage *versus* tirzepatide (Figure 6C, D).

**Figure 6.**
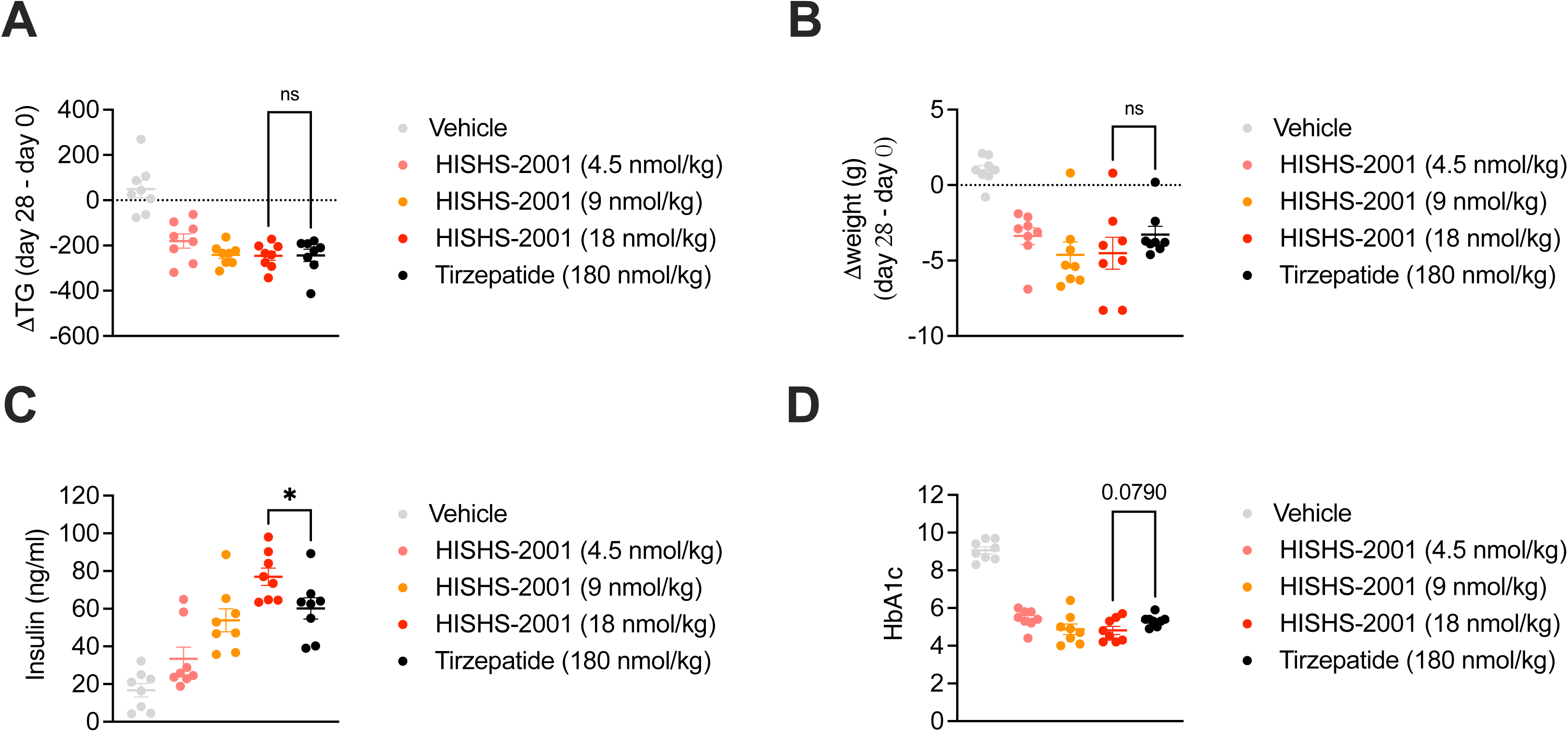
*In vivo* effects of HISHS-2001 *versus* tirzepatide in obese *db/db* mice. (**A**, **B**) Change (day 28 – day 0) in circulating triglycerides (**A**) and body weight (**B**) in obese mixed sex *db/db* mice chronically exposed to the indicated agonist; n=8. (**C**, **D**) Plasma insulin (C) and HbA1c (D) levels measured at day 28 in mice from (A, B). Data is shown as mean +/- SEM; ns, non-significant; *p<0.05, by one-way ANOVA with Sidak’s post-hoc test.

## Discussion

The main aim of the present study was to synthesize a novel GLP1R-GIPR agonist and perform its functional characterization *in vitro* and *in vivo*. We show that HISHS-2001 has several favourable characteristics, including a higher affinity at the GLP-1R and a more pronounced cAMP over β-arrestin 2 biased signaling profile when compared to tirzepatide. We also observed small apparent differences in the G protein coupling abilities of HISHS-2001 and tirzepatide at the GLP-1R and the GIPR. On the other hand, signaling towards intracellular Ca^2+^ changes, chiefly involving alterations in plasma membrane potential and Ca^2+^ influx though voltage-gated calcium channels^24^ with little or no mobilization via G*_α_*_q_-coupled receptors^25^, was not different between the two agonists. The changes in signaling bias described above were associated with only subtle differences in GLP-1R trafficking and cholesterol-rich nanodomain recruitment of the receptor in pancreatic beta cells, and no significant differences in the *ex vivo* capacity of these agonists to potentiate insulin secretion in mouse or human islets was detected.

We note that differences may exist between the contribution of GLP-1R and GIPR signaling for the effects of tirzepatide in mouse *versus* humans, and that the apparent advantages of GLP-1R/GIPR co-agonists in mice may simply reflect altered signaling properties (e.g, bias) at the GLP-1R^26^. Nevertheless, others have shown the importance of GIPR signaling for the effects of tirzepatide in mice, which exceed those of semaglutide^27^, and as such our use of a mouse model here was valid to demonstrate the non-inferiority of HISHS-2001 *versus* tirzepatide *in vivo*. Importantly, we provide validation for our findings using human islets, where responses to HISHS-2001 closely correlated with those observed in mouse islets. Future human-centric *in vivo* studies, involving the use of mice humanised for the GIPR, and ultimately studies in humans, beyond the scope of the present report, will be required to fully assess the effects of GLP-1R/GIPR co-agonists such as HISHS-2001.

Importantly, explored *in vivo* in a mouse model of obesity-related diabetes (db/db), HISHS-2001 was equally effective as tirzepatide in reducing triglyceride levels and body weight, even though the latter drug was used at a considerably higher dose. Moreover, HISHS-2001 was more effective than tirzepatide at increasing circulating insulin levels and tended to cause a more marked lowering in HbA1c, indicating that our *in vitro* findings can be extended to the *in vivo* setting. Although this could conceivably reflect differences in the pharmacokinetic profile of the two drugs, our preliminary measurements (Supplementary Table 2) failed to reveal a marked difference in this parameter. In any case, this explanation would seem unlikely given the higher dose of tirzepatide used in our study.

The effects of both drugs on body weight are likely, at least in a large part, to reflect a reduction in appetite and feeding (Supplementary Table 3). These changes are presumably the result of actions at feeding centres in the brain, which may include agouti-related protein (AgRP) neurons in the arcuate nucleus^28^ as well as other neuronal populations^29^. Central effects of GLP-1R (and GIPR) agonism may also be involved more directly (i.e. independently of weight loss, but involving changed neural inputs to the endocrine pancreas) in the actions of HISHS-2001 to increase insulin secretion^30,31^. We note that the contribution of GIPR agonism to the effects of HISHS-2001 and other co-receptor agonists *in vivo* remains to be resolved given that antagonism at this receptor also promotes weight loss^32^. However, differences in signaling bias between HISHS-2001 and tirzepatide at the GLP-1R, as demonstrated here in studies in model cell systems, do not appear to be important in central neurons in controlling body weight. Future studies, involving investigations of food intake and energy expenditure outside the scope of the present report will be needed to explore these questions in more detail.

## Conclusion

We suggest that HISHS-2001 provides an attractive dual receptor agonist which may be useful clinically as an antihyperglycemic agent in human obesity and T2D.

## Acknowledgements

The study was supported by a grant from Sun Pharmaceutical Industries to G.A.R. and A.T. The A.T. lab is funded by an MRC Project Grant (MR/X021467/1) and a Wellcome Trust Discovery Award (301619/Z/23/Z). AT also acknowledges funding from Diabetes UK, the Society for Endocrinology and the Eli Lilly LRAP programme. G.A.R. was supported by a Wellcome Trust Investigator Award (WT212625/Z/18/Z), MRC Programme grant (MR/R022259/1), Diabetes UK (BDA 16/0005485) and NIH-NIDDK (R01DK135268) project grants, a CIHR-JDRF Team grant (CIHR-IRSC TDP-186358 and JDRF 4-SRA-2023-1182-S-N), CRCHUM start-up funds, and an Innovation Canada John R. Evans Leader Award (CFI 42649).

## Contributor Information

Guy A. Rutter.

Alejandra Tomas

## Data availability statement

All data is available in the main article and/or the Supplementary Material.

