## Supplementary Information for "Binding Kinetics, Bias, Receptor Internalization and Effects on Insulin Secretion *in vitro* and *in vivo* of a Novel GLP-1R/GIPR Dual Agonist, HISHS-2001"

**Supplementary Table 1: Human islet preparations used in the study.** COD, cause of death; CVD, cardiovascular disease.

| Age (Years) | Sex | BMI (Kg/m$^2$) | HbA1c (%) | COD | Origin |
|---|---|---|---|---|---|
| 63 | M | 29.4 | - | CVD | Pisa |
| 76 | F | 23.9 | - | CVD | Pisa |
| 32 | M | 24.8 | 4.5 | - | Alberta |
| 47 | M | 40.3 | 5.5 | - | Alberta |
| 64 | F | 22.07 | - | CVD | Pisa |
| 53 | F | 27.2 | - | - | Milan |

**Supplementary Table 2: Pharmacokinetic analysis of HISHS-2001 *versus* tirzepatide.** See main manuscript text for further details.

| Treatment 30 nM/kg (N=5) | $AUC_{0-t}$ (hr*ng/mL) | $AUC_{0-\infty}$ (hr*ng/mL) | $C_{max}$ (ng/mL) | $T_{max}$ (hr) | $T_{1/2}$ (hr) | $K_{el}$ (1/hr) |
|---|---|---|---|---|---|---|
| **HISHS-2001** | 17307.8 | 17345.2 | 779.90 | 8.0 | 10.09 | 0.069 |
| **Tirzepatide** | 14080.6 | 14090.3 | 685.85 | 8.0 | 8.78 | 0.079 |

**Supplementary Table 3: Effects of HISHS-2001 *versus* tirzepatide on cumulative food intake.** See main manuscript text for further details.

| Treatment | Cage No. | Food intake (g), Day 28 (N=4) | Mean (g) | SD |
|---|---|---|---|---|
| | | | N=8 | |
| **Diabetic Control** | 1 | 212.4 | 202.5 | 14.00 |
| | 2 | 192.6 | | |
| **HISHS-2001 4.5 nM/kg (q3d*10)** | 3 | 116.85 | 112.4 | 6.28 |
| | 4 | 107.975 | | |

| | | | | |
|---|---|---|---|---|
| **HISHS-2001 9 nM/kg (q3d*10)** | 5 | 108.2 | 111.8 | 5.11 |
| | 6 | 115.425 | | |
| **HISHS-2001 18 nM/kg (q3d*10)** | 7 | 78.55 | 89.5 | 15.49 |
| | 8 | 100.45 | | |
| **Tirzepatide 180 nM/kg (q3d*10)** | 9 | 87.65 | 90.1 | 3.43 |
| | 10 | 92.5 | | |
